## Supplementary figures for "A novel RyR1 inhibitor prevents and rescues sudden death in a mouse model of malignant hyperthermia and heat stroke"

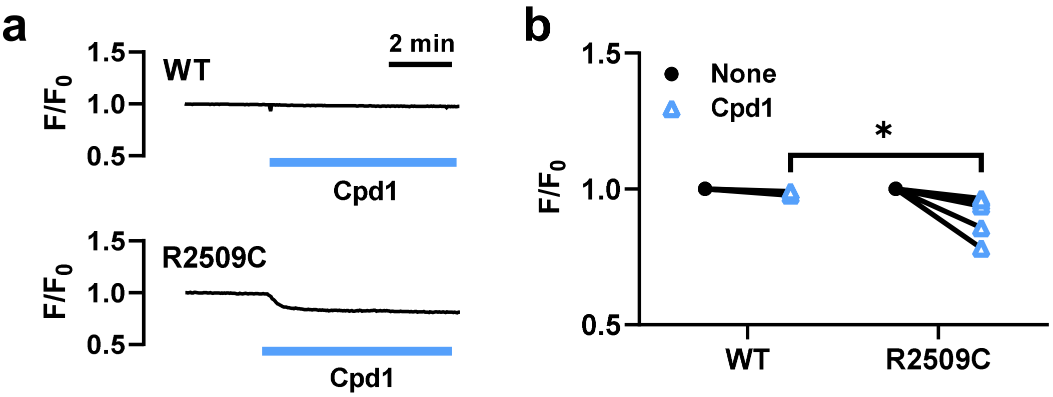


**Supplementary Figure 1. Effect of Cpd1 on resting $Ca^{2+}$ in the isolated FDB muscle cells from WT and R2509C mice. a.** Representative effects of 0.1 µM Cpd1 (blue bar) on resting $Ca^{2+}$ signals in FDB cells. **b**. Resting $Ca^{2+}$ levels before (circles) and during (triangles) application of 0.1 µM Cpd1. WT: n=5, R2509C: n=6. *p <0.05 compared to WT.

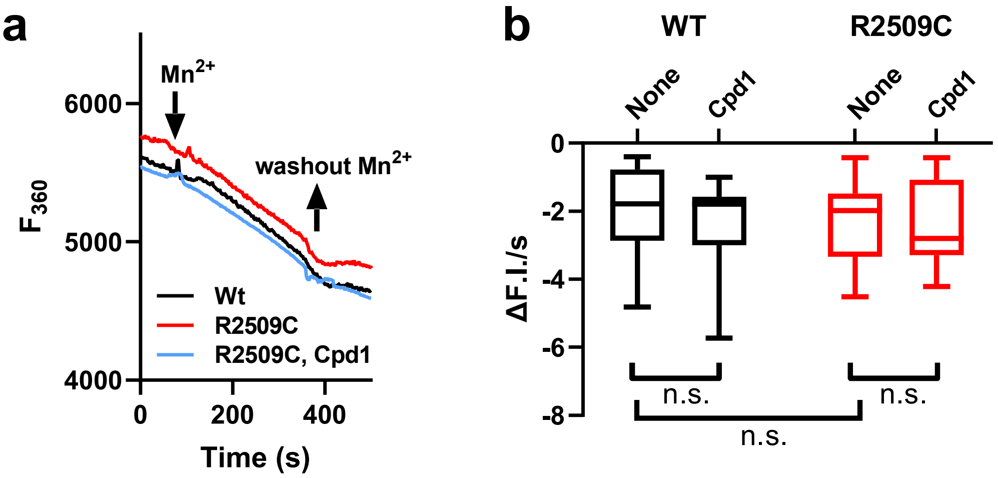


**Supplementary Figure 2. $Mn^{2+}$ quench assay with the isolated FDB muscle cells from WT and R2509C mice. a.** Representative fluorescence declines in fura-2 at 360 nm excitation by $Mn^{2+}$ in the external solution. Slopes for WT, R2509C and R2509C with 0.1 µM Cpd1 were very similar to each other. **b.** Average decline rates of fura-2 fluorescence in the absence and presence of 0.1 µM Cpd1. Data are expressed as box-whisker plots (WT: n=16; WT, Cpd1: n=14; R2509C: n=18; R2509C, Cpd1: n=12). n.s., not significant.

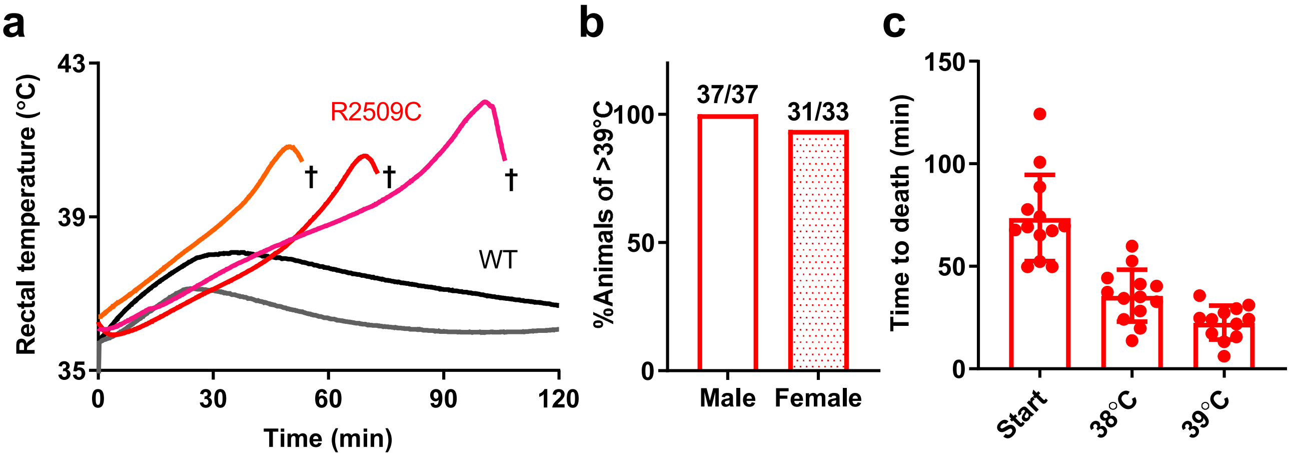


**Supplementary Figure 3. *In vivo* heat stress challenge of WT and R2509C mice.** Mice were anesthetized and placed in a test chamber at 35°C. **a.** Rectal temperature of mice. R2509C mice but not WT mice exhibited rise in rectal temperature and died by fulminant heat stroke (†). **b.** Responsiveness to heat stress. Almost all the mice responded to heat stress. **c.** Time to death from start of heat stress challenge or from temperature at 38°C or 39°C. Data are means±SD (Start: n=13; 38°C: n=13; 39°C: n=12).

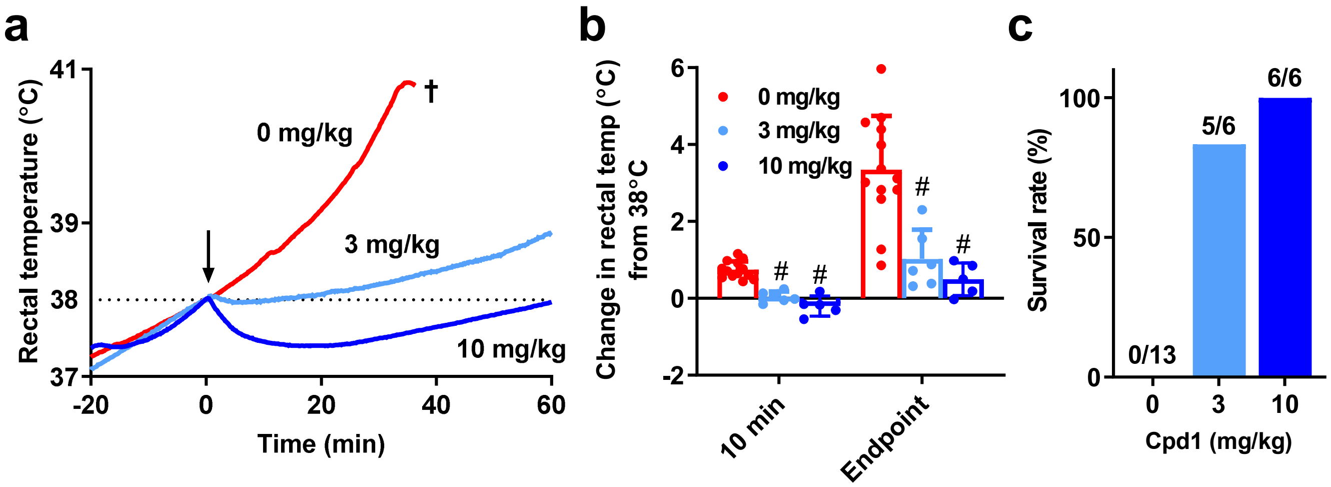


**Supplementary Figure 4. Rescue effect of Cpd1 on heat stress challenge in R2509C mice.** Cpd1 (0, 3, or 10 mg/kg) was administered *i.p.* when their body temperature reached 38°C. **a.** Rectal temperature in R2509C mice after administration of Cpd1 (arrow) during heat stress challenge. †, death by heat stroke. **b.** Change in the rectal temperature 10 min after administration of Cpd1 and the endpoint (60 min after administration or just before death). Data are means±SD (0 mg/kg: n=13; 3 mg/kg: n=6; 10 mg/kg: n=6). #p <0.05 compared to 0 mg/kg. **c.** Survival rate of mice 60 min after administration of Cpd1.

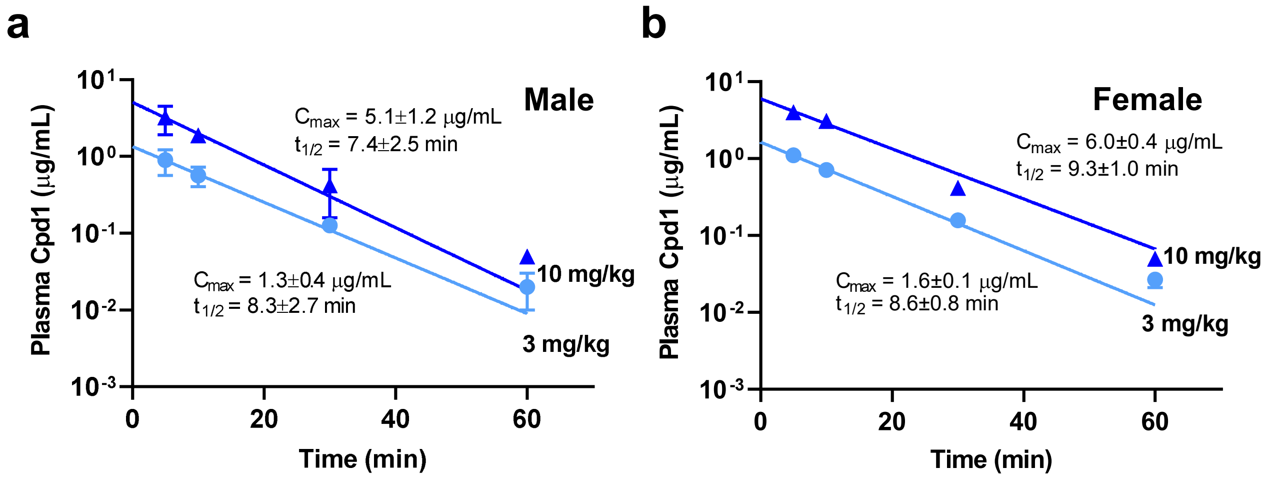


**Supplementary Figure 5. Pharmacokinetics of Cpd1 in mice**. Average plasma concentration-time profiles of Cpd1 following *i.p.* injection of 3 mg/kg and 10 mg/kg in male (**a**) and female (**b**) mice. Data are mean±SD (n=3 per time point).

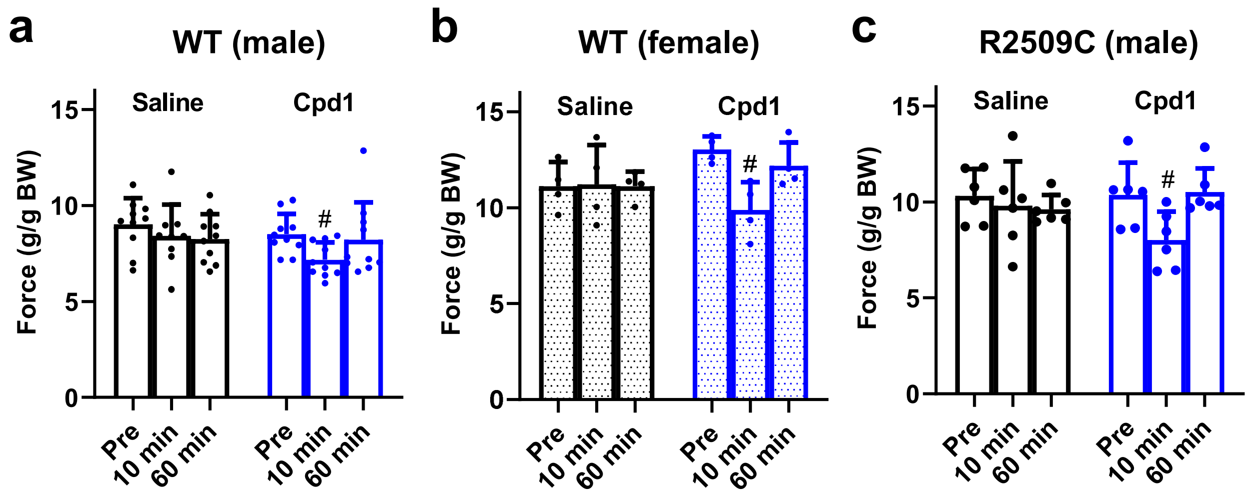


**Supplementary Figure 6. Effect of Cpd1 on muscle force of WT and R2509C mice.** Muscle force of male WT (**a**), female WT (**b**) and male R2509C (**c**) mice was measured *in vivo* by grip force test (4-grips test). The tests were performed before (*pre*) and 10 and 60 min after *i.p.* injection of saline or Cpd1. The force values were normalized by body weight (BW). Data are mean±SD (male WT: n=10; female WT: n=4; male R2509C: n=6). #$p$<0.05, compared to the value before injection (*Pre*).

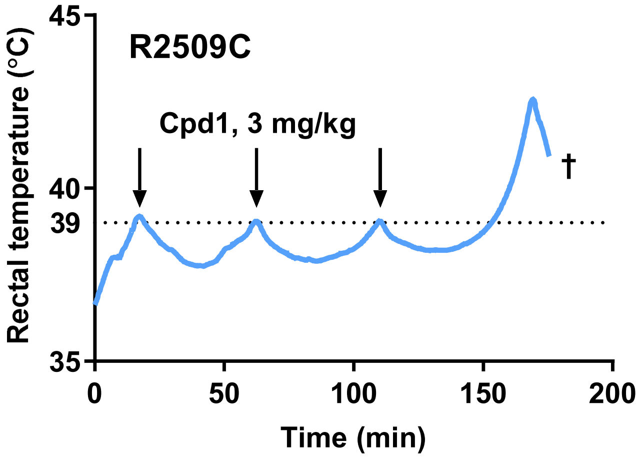


**Supplementary Figure 7. Effect of repeated application of Cpd1 on *in vivo* isoflurane challenge of R2509C mouse.** Rectal temperature in male R2509C mouse was measured after anesthesia by isoflurane. Cpd1 (3 mg/kg) was repeatedly administered *i.p.* when the body temperature reached 39°C (arrow). †, death by fulminant MH crisis.

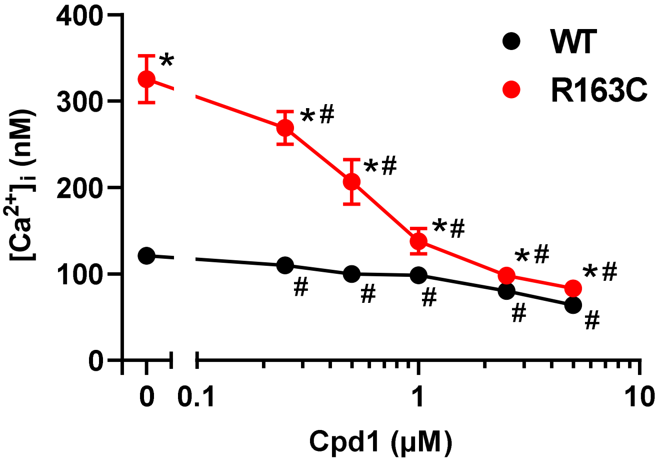


**Supplementary Figure 8. *In vitro* $[Ca^{2+}]_i$ homeostasis of skeletal muscles isolated from WT and R163C mice.** FDB muscle fibers were isolated and $[Ca^{2+}]_i$ was determined using $Ca^{2+}$ selective microelectrodes. $[Ca^{2+}]_i$ was significantly higher in quiescent R163C fibers (326±27 nM, =16) compared to WT fibers (121±3 nM, n=20). Cpd1 reduced $[Ca^{2+}]_i$ in R163C fibers in a dose-dependent manner. Data are means±SD (n=10-20). *p<0.05 compared to WT. #p <0.05 compared to the value without Cpd1.
